# The power of peer networking for improving STEM faculty job applications: a successful pilot program

**DOI:** 10.1101/2021.10.16.464662

**Authors:** Carlos M. Guardia, Erin Kane, Alison G. Tebo, Anna A.W.M. Sanders, Devrim Kaya, Kathleen E. Grogan

## Abstract

In order to successfully obtain a faculty position, postdoctoral fellows or ‘postdocs’, must submit an application which requires considerable time and effort to produce. These job applications are often reviewed by mentors and colleagues, but rarely are postdocs offered the opportunity to solicit feedback multiple times from reviewers with the same breadth of expertise often found on an academic search committee. To address this gap, this manuscript describes an international peer reviewing program for small groups of postdocs with a broad range of expertise to reciprocally and iteratively provide feedback to each other on their application materials. Over 145 postdocs have participated, often multiple times, over three years. A survey of participants in this program revealed that nearly all participants would recommend participation in such a program to other faculty applicants. Furthermore, this program was more likely to attract participants who struggled to find mentoring and support elsewhere, either because they changed fields or because of their identity as a woman or member of an underrepresented population in STEM. Participation in programs like this one could provide early career academics like postdocs with a diverse and supportive community of peer mentors during the difficult search for a faculty position. Such psychosocial support and encouragement has been shown to prevent attrition of individuals from these populations and programs like this one target the largest ‘leak’ in the pipeline, that of postdoc to faculty. Implementation of similar peer reviewing programs by universities or professional scientific societies could provide a valuable mechanism of support and increased chances of success for early-career academics in their search for independence.

## Introduction

The purpose of a postdoctoral appointment is the acquisition of additional skills or training post-PhD in preparation for transitioning into an independent position as a primary scientific investigator (National Research Council 2014), originally in academia (NAS et al. 2000), although the function has broadened to include training for a multitude of other career paths (McConnell et al. 2018). Thus, by definition postdoctoral appointments are temporary (average = ∼2.7 years; Acton et al. 2019, Andalib et al. 2018, McConnell et al. 2018, but see Fernandes et al. 2020), although individuals may complete multiple postdoctoral positions in different labs before gaining independence (Powell 2020, Shaw et al. 2015, Woolston 2020). Because of the limited duration and lack of funding security for postdoctoral positions, and the highly competitive tenure-track faculty job market (Alund et al. 2020, Andalib et al. 2018, Kahn & Ginther 2017, Sauermann & Roach 2012, Zimmerman 2018), postdocs spend a significant amount of time searching for their next appointment. Some will begin to search for the next position as soon as they begin their current position and many apply for postdoctoral and tenure-track positions simultaneously over multiple years (Fernandes et al. 2020).

An application packet for a faculty position (‘faculty application’) consists of multiple highly crafted documents, typically including a curriculum vitae (CV), cover letter, statements of research, teaching, and, sometimes, diversity (for a description of these documents and the faculty application process, see Fernandes et al. 2020, Groll 2017). To maximize the chance of success, applicants spend a significant amount of time writing and polishing these documents. Numerous opinion and advice pieces have been published on how to write these documents (e.g., Anderson 2019, Reyes 2020, Smith 2020a & 2020b). In addition, Offices of Postdoctoral Education across institutions, as well as the National Postdoctoral Association in the USA, and some scientific societies host frequent seminars/webinars and provide extensive advice on the structure and content of these documents (e.g., Omary et al. 2019, Shaw et al. 2015). Almost all of these seminars and advice columns direct postdocs to solicit feedback from a wide circle of peers and mentors.

Formal and informal mentors are often willing to provide constructive feedback (Hayter & Parker 2019); however, some mentors may be unwilling to spend time on this task or unable to offer useful feedback, especially for postdocs applying for positions in fields or institutions different than the mentors’ own (Alund et al. 2020, Aschwanden 2006, Hayter & Parker 2019, Scaffidi & Berman 2011). Postdoctoral peers and senior graduate students can also serve as additional reviewers, but most research groups only have a couple of postdocs at a time or may have none at all (Acton et al. 2019, Bruckman & Sebestyen 2017). Moreover, the breadth of scientific expertise represented within a research group or postdoc’s network rarely matches the breadth of expertise represented by search committees in academia. Thus, while one’s colleagues/labmates may be able to comment on the structure and the science within a job application, they may not be able to assess if a research statement is broad and general enough to be understood by, and appeal to the wider audience represented by a search committee. Furthermore, while mentors and peers may be happy to review a document a few times, most mentors and peers lack the time to provide multiple rounds of feedback on >10 pages of job application materials.

Overall, opportunities to have job applications critiqued repeatedly by a broad scientific audience are generally scarce. Regardless of the underlying causes, postdocs could benefit from a variety of options for getting feedback on their job application materials. This situation is exacerbated for postdocs from marginalized groups, who are more likely to struggle to form a supportive peer network and have less access to mentoring and support than postdocs from majority groups (Beech et al. 2013, Yadav & Seals 2019). Therefore, here, we present a potential solution in the form of an open and inclusive international program of peer review for job application materials that we have been running since 2018. In this program, participants have the opportunity to repeatedly share all or part of their application package with a small group of peer reviewers. They engage in reciprocal constructive commentary in a supportive and encouraging manner with the ultimate desire of seeing each other attain an independent faculty position. Peer review programs are most well-known from their use in manuscripts (Rennie 2016, Tennant et al. 2017, Tennant & Ross-Hellauer 2020, but see Haffer et al. 2019, Murray et al. 2018) and grant application review (Azoulay & Li 2020, Demicheli et al. 2007, Marsh et al. 2009 but see Lauer & Roychowdhury 2021, Witteman et al. 2019), but have been successfully implemented in many other contexts for purposes of professional development and community building, especially for postdocs and early career researchers (Dickson et al. 2021, Eisen & Eaton 2017, Kulage et al. 2017).

This manuscript has three goals: i) to describe the history and organizational details of our program; ii) to use survey data to assess the experiences of participants and nonparticipants on the benefits and limitations of this type of program; and iii) to suggest methods of implementation for other organizations (i.e., Offices of Postdoctoral Education, postdoctoral societies, or scientific societies).

### Program Description

This program began organically after its founder, Dr. Grogan, observed frequent requests for peer review of job application materials on the FuturePI Slack Group (https://futurepislack.wordpress.com/) and realized the group could benefit from the organization of formal peer-reviewing of application materials during the Fall 2018 job application season. In its first year, the FuturePI Reviewing Groups program (hereafter called the Program) ran for 7 weeks, from mid-August to mid-October. In subsequent years, the timeline expanded to 15 weeks, from early August to the end of November (Supplemental Material). The program is announced through the FuturePI Slack #general and #Jobapp_reviewer channels two weeks before its start to give participants time to sign up, with sign-ups handled on an open-access Google Sheet. Participants are asked to provide their names, email addresses, general field of study, and the type of jobs they are applying to, and to indicate which weeks they would like to participate (see Supplemental Figure 2 for example). Reviewing groups are organized weekly, with sign-ups for the upcoming week closing on Sunday morning. All interested participants for a given week are emailed the day before groups are assigned to confirm their willingness to participate that week, and then reviewing groups are organized the next morning. Each reviewing group is emailed at the start of the week with contact information for their group and instructed to send whatever documents they want to be reviewed and to provide feedback on each other’s documents by the end of the week (for example announcement, confirmation, and assignment emails, see Supplemental Materials). For the first three years of operation, participants were instructed to send their documents to each other on Monday and provide feedback for group members’ documents by Friday, but this schedule can be adjusted easily. Participation in the program is open to any current member of FuturePI Slack and the program has grown since its first year, from 21 unique participants in 2018 to 71 in 2020 (Figure 1A). The number of unique individuals who participate each week varies considerably (mean = 8.7, range = 3-17; Figure 1B), but has steadily increased since the program’s beginning (F = 4.363, p = 0.02, Figure 1C). Additionally, the number of times that any given individual participates has also increased, although not significantly (F = 1.039, p = 0.36; Figure 1D).

**Figure 1.**
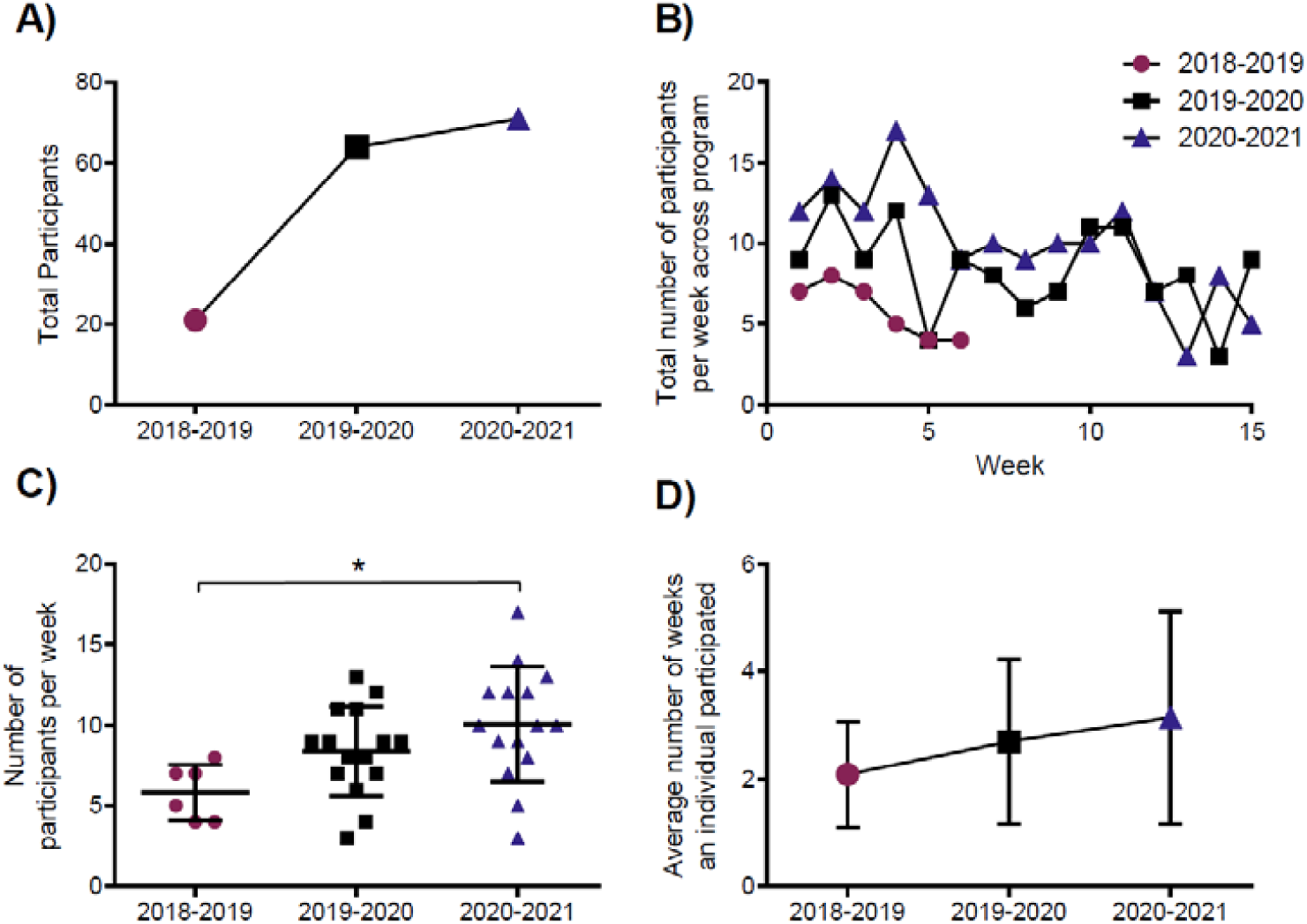
Program participation for the FuturePI Reviewing Groups Program from 2018 to 2021. A) The total number of unique individuals who participated in the reviewing program per year, B) the total number and C) average of individuals who participated each week of the program by year, D) the average number of weeks an individual participated by year.

**Figure 2.**
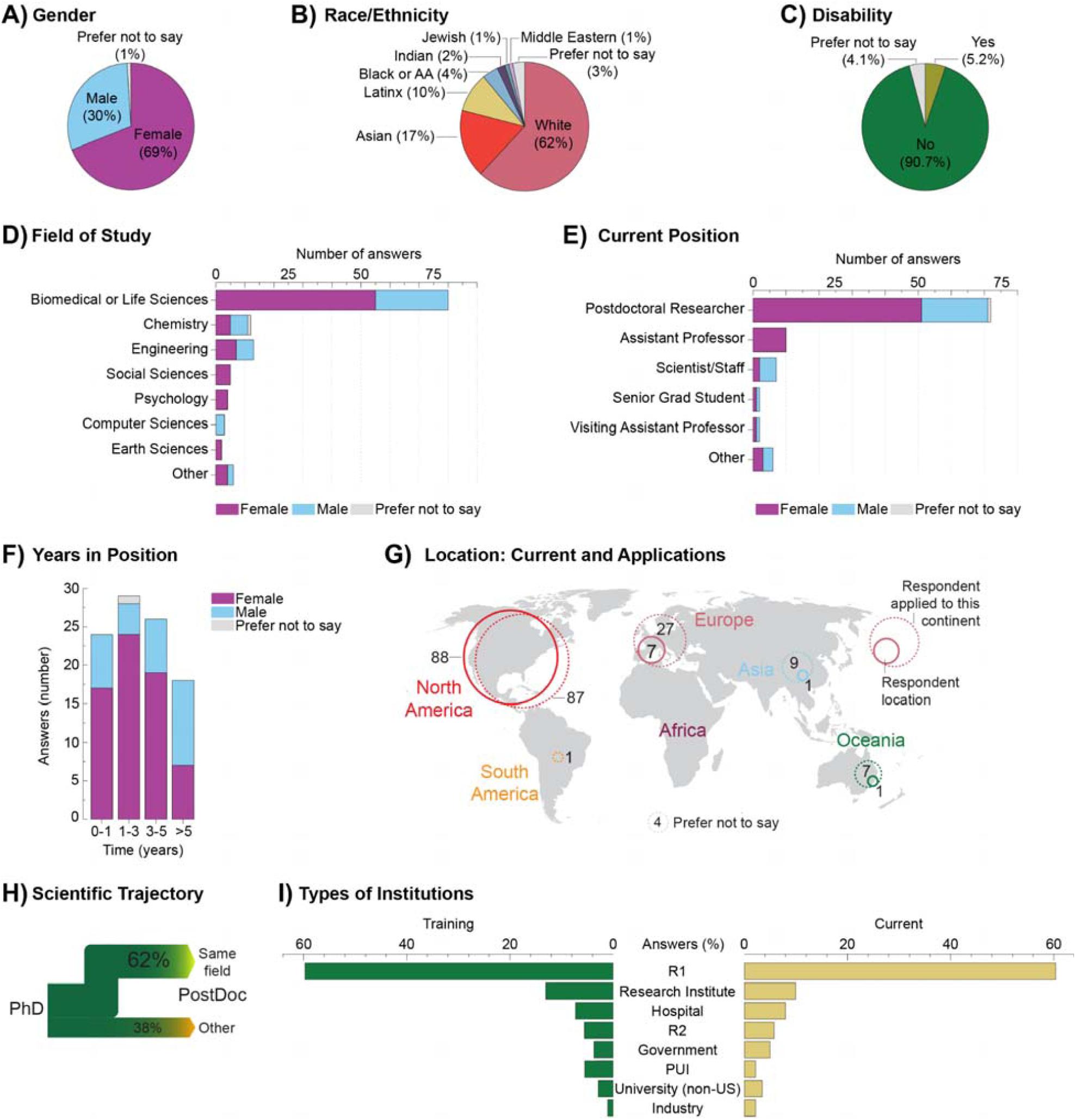
Self-identified demographics and career information of survey respondents, which included both program participants and non-participants: the distribution of survey respondents by A) gender, B) race/ethnicity, and C) disability status; The distribution of respondents, split by gender, according to D) field of study, E) current position title, and F) years in current position; G) map of the current location of survey respondents (solid circles) compared to the location of positions to which respondents applied (dashed circles); H) the representation of percentage of survey respondents who either remained in the same field as their Ph.D. training or switched scientific fields post-graduation; I) the type of institution at which survey respondents were or are employed.

### Program Feedback Survey

Given the increase in participation and anecdotal commentary about the benefits of participating in the program, we developed a survey for participants and non-participants in the Program to assess participants’ experiences and identify areas for improvement (Supplemental Table 1). Through this survey (University of Cincinnati IRB# 2020-0891), we collected demographic and participation-related information from members of the FuturePI Slack community. To maintain anonymity and promote ease of response, the survey, which took ∼5-10 minutes to complete, was conducted through Google Forms and all questions were optional, except the IRB permission. Respondents were recruited from previous and current participants in the program either through messages posted on FuturePI Slack or through direct email. Responses were collected within two months starting in early February and ending at the end of March 2021. Data were organized and analyzed in Excel, OriginPro2015, and GraphPad Prism 6 respectively. Answers to open-ended questions were stripped of white space, punctuation, numbers, and English “stop words” (e.g., and, the, is) using the package *tm* (Version 0.7-8; Feinerer & Hornik 2020) and after manual editing to change plural nouns to singular nouns and stem words with the same root (e.g., standardizing review, reviews, and reviewed as “review”). Then word clouds were generated using the package *wordcloud* (Version 2.6; Fellows 2018) in RStudio.

#### Survey Respondent Demographics

All individuals who participated in the survey, regardless of whether they participated in the Program, are hereafter referred to as respondents, while respondents who participated in the Program are referred to as participants and those who did not participate as non-participants, respectively. The majority of our survey respondents (n = 97), which included both program participants and non-participants, identified as female (69%), white (62%), and did not identify as disabled (90.7%; Figure 2A-C). The respondents of this study were more likely to identify as female compared to the respondents of the 2016 USA National Postdoctoral Survey and a recent survey of faculty job applicants, in which 53% and 48.2% of respondents identified as female, respectively (Fernandes et al. 2020, McConnell et al. 2018, but see Afonja et al. 2021). In contrast, in this study, the self-identified race/ethnicity of the respondents was similar to that reported for other postdoctoral or faculty application populations for white, Black or African American, and Latinx/Hispanic identities. Additionally, this study unexpectedly included fewer Asian respondents (17.0%) compared to the percentage of Asian postdocs in the US postdoctoral population (24.8%; McConnell et al. 2018). Nearly 10% of our respondents identified as having a disability, higher than the 6% (12 out of 175) reported by a similar survey of postdocs and early career researchers in ecology and evolution (Wanelik et al. 2020).

Compared to a recent survey of faculty applicants (also organized in part through FuturePI Slack; Fernandes et al. 2020), the respondent population of this study was similarly biased towards postdocs over graduate students, staff scientists, or Assistant Professor and had many more respondents working in Biomedical or Life Sciences, with less than 15 respondents each from Chemistry, Engineering, Social Sciences, and other fields (Figure 2D-E). As FuturePI Slack is an international organization, the survey respondents were located across the globe and were applying to a similarly broad geographic range of jobs (Figure 2G). We had a nearly equal number of survey respondents across the various ranges of job tenure from 0-5 years. Nearly 54% of respondents were currently employed at an R1 institution (Carnegie classification), with the rest of the respondents mostly employed at research institutes, hospitals, R2 or PUI institutions, or in government positions (Figure 2I). Fewer respondents (40.8%) reported being trained at an R1 institution compared to other types of institutions. Interestingly, nearly 40% of our survey respondents reported they had changed fields between their PhD and their current position (Figure 2H). Of the 97 survey respondents, a majority (n = 58; Figure 5A) were participants in the Program at least once in the previous three years while 39 respondents were non-participants.

#### Survey Respondent Job Application Journey

To assess the job application trajectory difference between participants and non-participants, the survey included questions detailing the number of years respondents had been on the job market, as well as the number of job applications submitted either prior to the program (participants) or prior to this year (non-participants). Respondents in their first year of the job market were the largest portion (>40% of total) for both participants and non-participants (Figure 3A). However, more than 30% of participants were in their third or greater year of the job search compared with only ∼15% of non-participants (Figure 3A, left two columns). Notably, three participants were in their sixth year on the job market. While the majority of respondents (∼80%) had submitted ten or fewer applications, some of the respondents had already submitted substantial numbers of applications, including six individuals who reported submitting >90 applications. In comparison, none of the non-participant had submitted more than 50 applications at the time of the survey. From these data, we observed that the participants tended to be further along in their job search trajectory, having spent more years searching for jobs and submitting many more applications.

**Figure 3.**
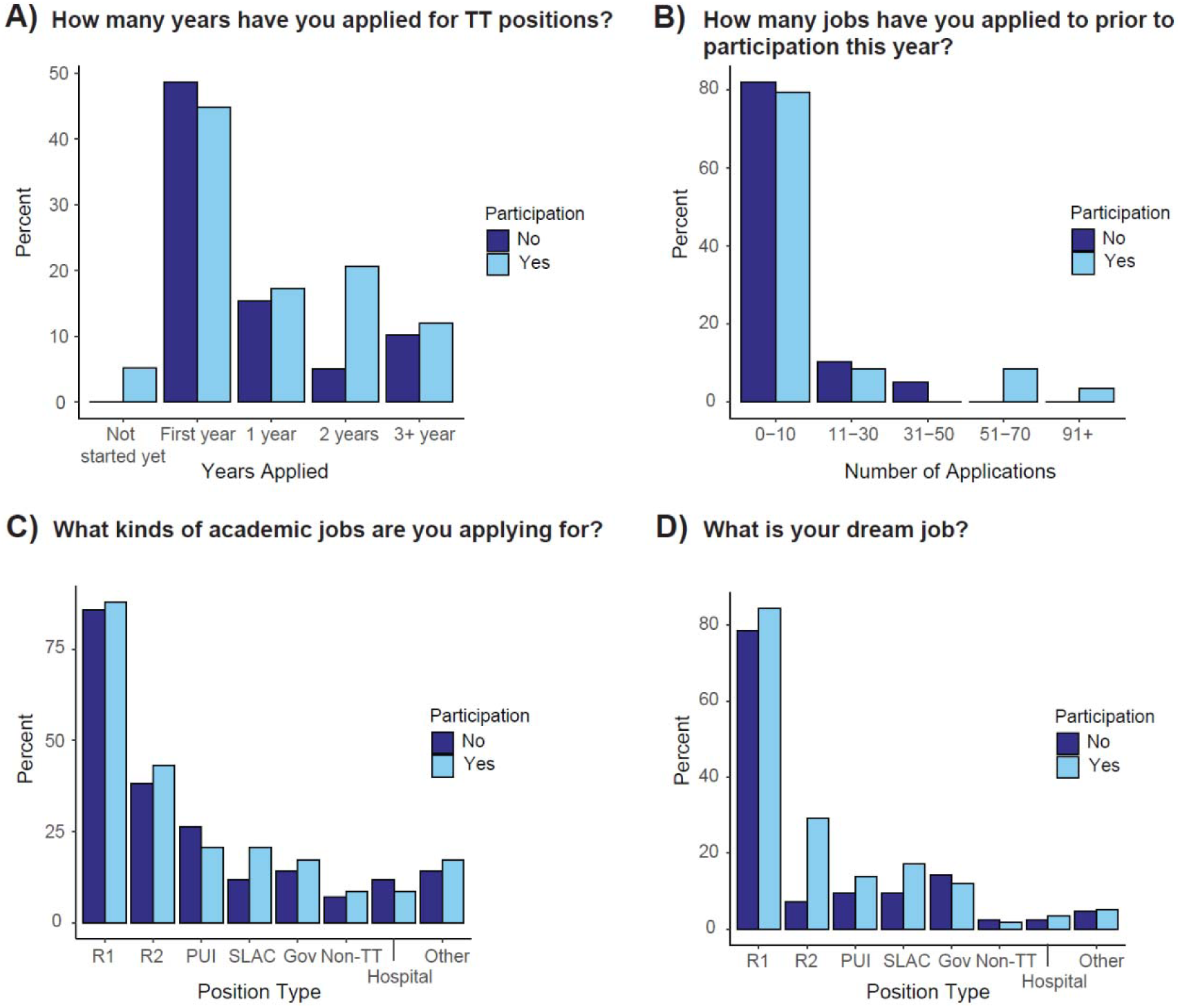
Comparison of the faculty job application journey for participants and nonparticipants: A) the number of years spent searching for a faculty job, B) number of job applications and C) types of job applications submitted by participants and nonparticipants, and D) the dream job for participants and nonparticipants. For C & D, Other includes applications for faculty, research scientist, and postdoctoral positions at Professional Schools, Research Institutes, non-profits, and industry companies

**Figure 4.**
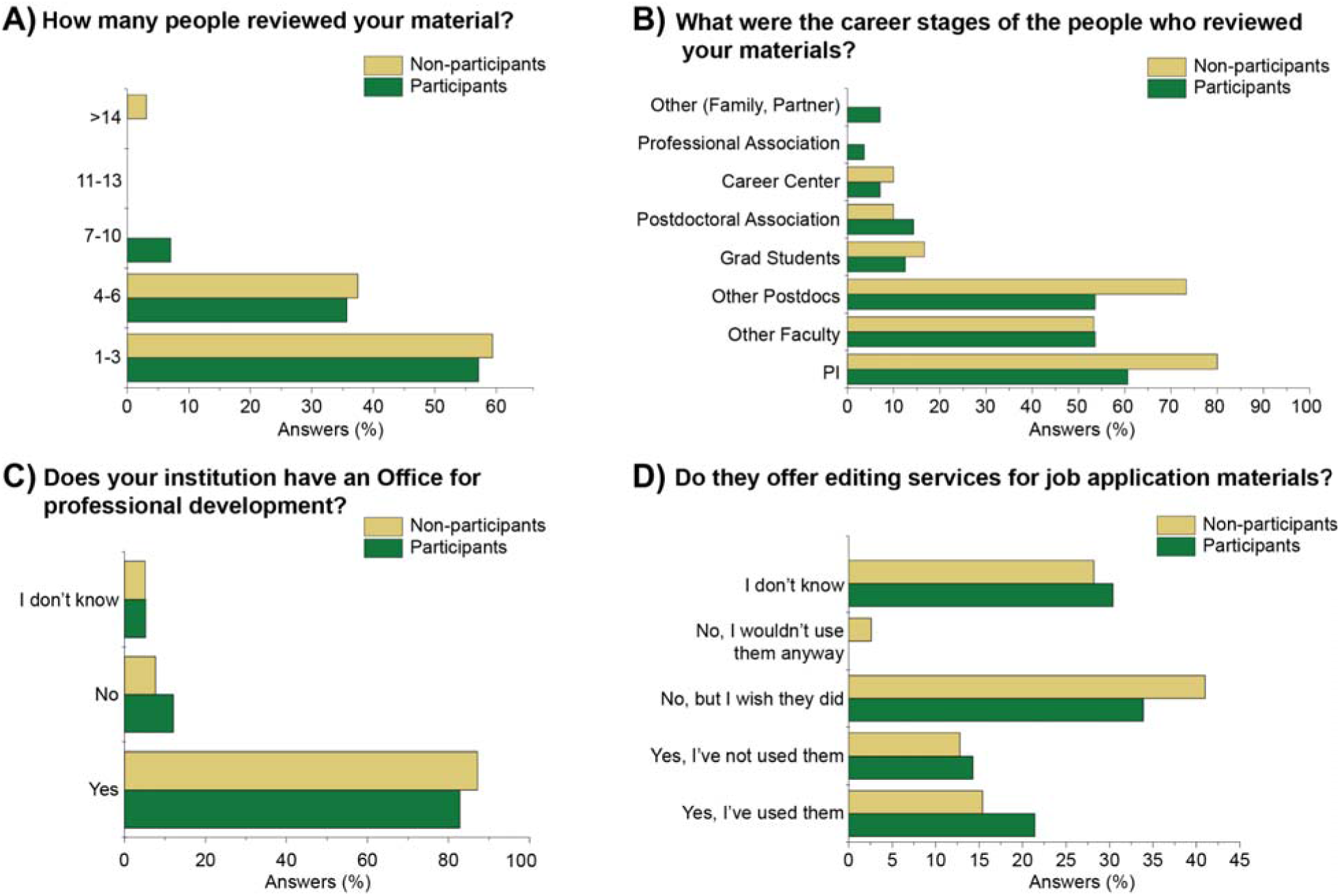
Information on the availability of reviewers for survey respondents’ job application materials: A) the number and B) career stages of reviewers recruited by survey respondents to review their job application materials, C) the percentage of survey respondents who work at an institution that has an Office focused on professional development, D) if those offices offer job application editing services, and if the respondents have taken advantage of those services or not.

The types of institutions that the respondents applied to as well as their ideal position were also examined in the survey. Among all respondents, R1 universities were the most applied to position (n = 51 participants and 33 non-participants of 97 total respondents) and most frequently reported dream position (n = 49 participants and 31 non-participants). The participants applied to more non-R1 positions compared to non-participants, which could be due to the career stage and experience of the participant in previous job cycles leading to a broadening of their targeted institution. More participants also reported that their dream jobs were at non-R1 educational institutions, e.g. R2 universities, primary undergraduate institutions (PUIs), and small liberal arts colleges (SLACs).

#### Reviewer Availability for Survey Respondent Application Materials

Most of the respondents had their application materials reviewed by either 1-3 people (55-60%) or 4-6 people (35-40%) outside the Program. The application material reviewers consisted of mostly PI, other faculty, and postdoctoral colleagues, but the percentage differed between participants and non-participants. For participants, slightly over 60% had their PI review their materials, ∼53% had other faculty reviewers, and nearly 55% recruited other postdoctoral colleagues to review their materials. In contrast, for non-participants, ∼80% were able to have their PI review their materials, ∼53% had other faculty reviewers, and nearly 75% were able to recruit other postdoctoral colleagues to review their materials. In other words, participants were less likely to have their job application materials reviewed by their PI or other postdoctoral colleagues but were equally likely to have non-PI faculty review their materials. Fewer than 15% of survey respondents, regardless of Program participation, asked graduate students, their postdoctoral association or career center, or other individuals to review their materials.

Although the majority of survey respondents (>80%) were employed at institutions with Offices for Professional Development, many respondents were either unsure (25-30%) if these offices offered editing services or reported these offices did not offer those services (∼35-45%). Of those respondents with professional development offices that offered editing services, the participants (22%) were slightly more likely to have used those services than non-participants (15%). Altogether, over 50% of both participants and non-participants indicated either a willingness to use editing services provided by offices of professional development or a wish that they would offer those services.

#### Program Participation Data

With nearly all participants indicating they had submitted the research statement for peer review, the research statement was the most reviewed document (Figure 5A) followed closely by the teaching statement, CV, and Cover Letter. The least reviewed document was the Diversity Statement, which may reflect that fewer job applications require this document or the more personal nature of this particular document. The majority (∼65%) of the participants reviewed documents and had their documents reviewed by 1-5 people (Figure 5B), implying they participated for 1-2 weeks. Receiving general feedback and seeing other job application examples were reported as the two most valuable aspects of participation by over 80% of the participants (Figure 5C). Refining materials, connecting other postdocs, and having an early deadline were rated as comparably less important or valuable aspects of participating in the program. Comments on the content were reported as most valuable and copy-editing being the least valuable type of feedback (Figure 5D). Participants reported extremely high satisfaction with their participation in the Program, with only 8.6% (5/58) giving a rating lower than 5 out of 7 (Figure 5E). Similarly, 96.5% of respondents (56/58) reported they would likely or highly likely to recommend participation in the Program to other colleagues.

**Figure 5.**
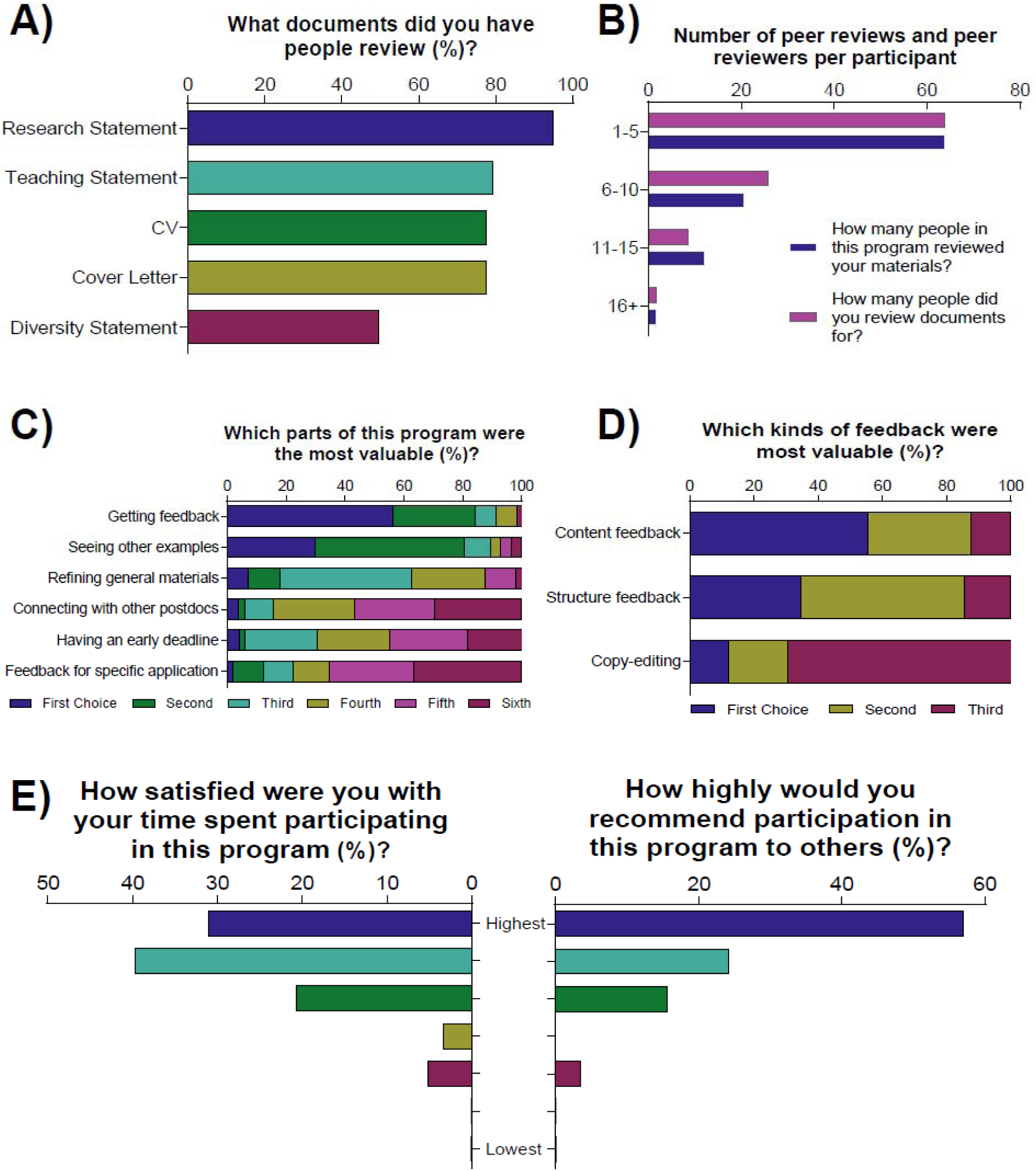
Participants were asked to indicate: A) the documents they had peer-reviewed in the program, and B) how many people reviewed the documents for each participant and how many people each participant reviewed documents for, C) the parts of the program and D) types of feedback that participants felt were most valuable, and E) participants indicated how satisfied they were with their participation in the FuturePI Reviewing Groups Program and how highly they would recommend participation in the program to others.

When asked about their motivations for participating in the Program, the respondents were primarily motivated (Figure 6A) to i) seek feedback beyond their own lab (n = 34), especially from peers (n = 12) and people outside of their subdiscipline (n = 16) and ii) to use examples from others to better organize their job documents (n = 10):

**Figure 6.**
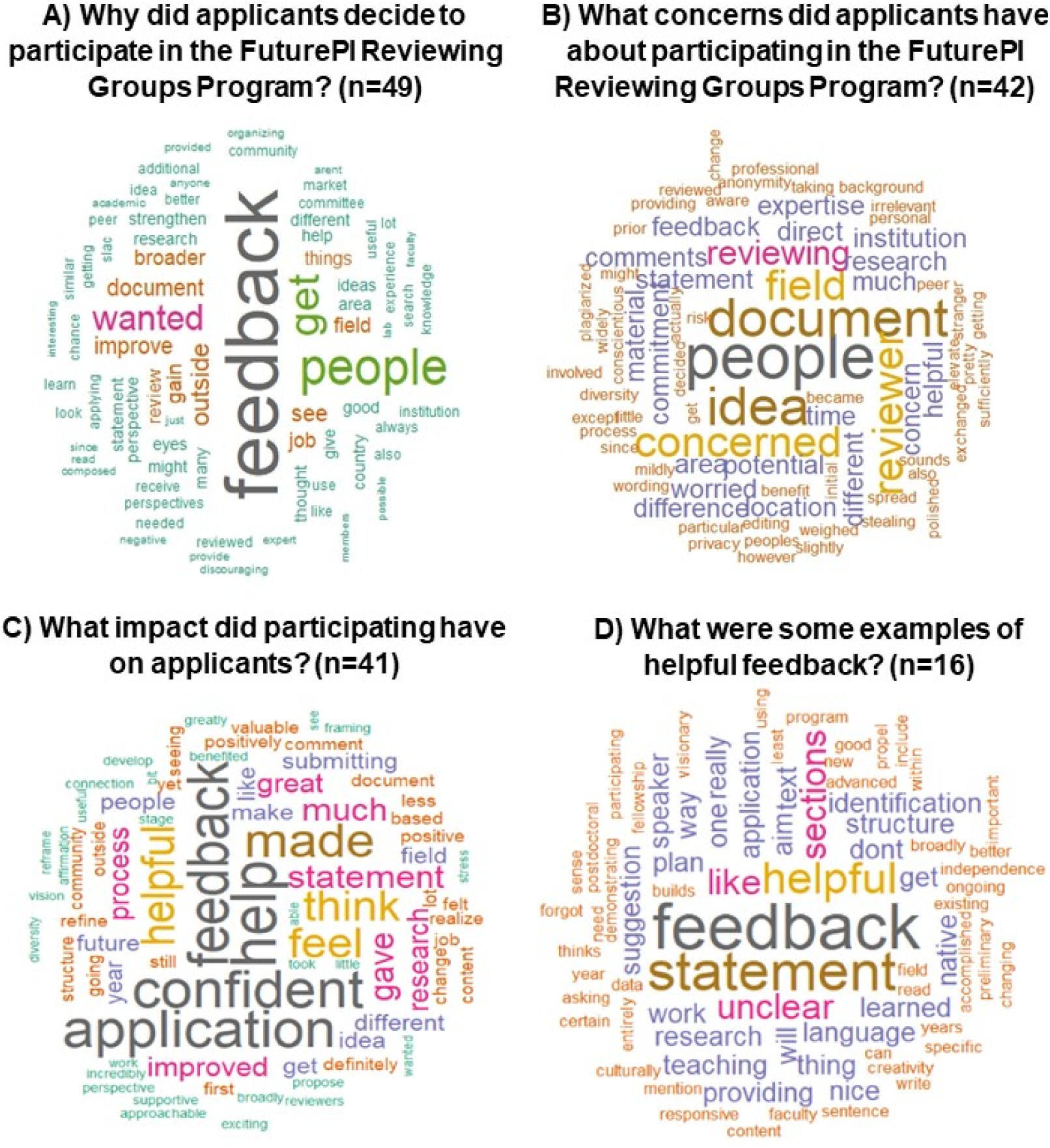
Word clouds highlighting the most common responses of the participants to the open-ended questions on: A) why they decided to participate in the program, B) what concerns they had about participating prior to doing so, C) what impact did participation have on their job application journey, and D) what were some examples of helpful feedback.

> *“I wanted a broader perspective than could be provided from just the members of my lab, since a faculty search committee will be composed of many people who aren’t experts in my area. I also wanted as many eyes as possible to read my documents. And I wanted to see the documents of others to see if anyone had any interesting ideas I could use*.*”*
>
> *“To get more positive feedback (my PI gave me very negative feedback and it’s [sic] discouraging)*.*”*
>
> *“It was an amazing opportunity to receive feedback from people unfamiliar with my work/area/research, and who could be honest with their feedback as they had no investment in the outcome*.*”*

The majority of participants noted, as shown in the representative quotes below, that their participation in the Program improved their confidence in their job application materials, the application process (n = 13; Figure 6C), and the quality of their materials (n = 17), and that they felt more supported during the job application process, which is often a stressful and isolating (n = 14):

> *“A little bit of positive affirmation took some stress out of the process*.*”*
>
> *“It gave me more confidence, both because someone had looked at my stuff and because I felt like a part of a community and that I didn’t have to do this all on my own - there were other great people like me going through the same thing*.*”*
>
> *“The first year was very helpful. As a first gen with a non-helpful PI, I had no idea of how the process worked or what was expected. The second year was less helpful as I felt participants were expecting solid feedback without putting in as much effort reviewing for others*.*”*
>
> *“I’m still on the job market, but it helped me realize that I’m in the same boat as everyone else*.*”*

In contrast, only three participants felt that their participation had little or no impact. When asked to provide examples of helpful feedback, the participants noted they received helpful feedback about the grammar and structure of their documents (n = 9) and identifying areas which needed more clarity (n = 8):

> *“One of my reviewers suggested a way to structure my teaching statement that I hadn’t considered. I still use it!”*
>
> *“Pointing out sections that would be unclear to a broader audience, suggestions on how to highlight future plans (i*.*e*., *what will your lab look like/do)”*
>
> *“Grammatical errors were plenty in my text, since I am not a native speaker. So it was very helpful for me that (apparently) a native English speaker edited my texts*.*”*
>
> *“Someone told me “This is really cool and I can’t wait to see what happens when you get a job!” which was so nice and affirming to hear - this whole process is so demoralizing and kind of dehumanizing, and it’s really hard to get feedback (and praise, honestly) from people who don’t already know you. This person offered helpful constructively critical feedback too, but that was such a nice boost*.*”*

The majority of participants, 29 out of 42, had no concerns about participating (69%). When the participants expressed concern, their concerns revolved primarily around competition and plagiarism (n = 8) or the lack of relevant feedback from reviewers (n = 7):

> *“In editing peers’ diversity statements in particular, I became a little concerned that this process might elevate people’s statements to sound more aware and conscientious than they actually are on their own*.*”*
>
> *“Some concerns about competition with those who may be applying to the same positions as myself*.*”*
>
> *“I was worried that my reviewers [sic] fields would be very different from my own*.*”*
>
> *“Some of the feedback I received seemed rushed and as though the other participant hadn’t put much effort into reviewing my documents*.*”*

Among the non-participants, 22 out of 38 were aware of the Program’s existence before the survey (Figure 7A), but many of these respondents either were not on the job market in the prior year (n = 6), were too busy (n = 6), or had gotten feedback elsewhere (n = 8). Of those that planned to go on the job market soon, most (11/19) planned to participate in the Program in the future or were undecided (6/19) - very few non-participants planned never to participate in the Program (2/19; Figure 7B). In our open-ended question about their reasons for not participating, non-participants reported that they generally heard about the Program too late or did not have job materials ready (n = 9), or they wanted feedback from different people than they thought participated (n = 6; Figure 7C):

**Figure 7.**
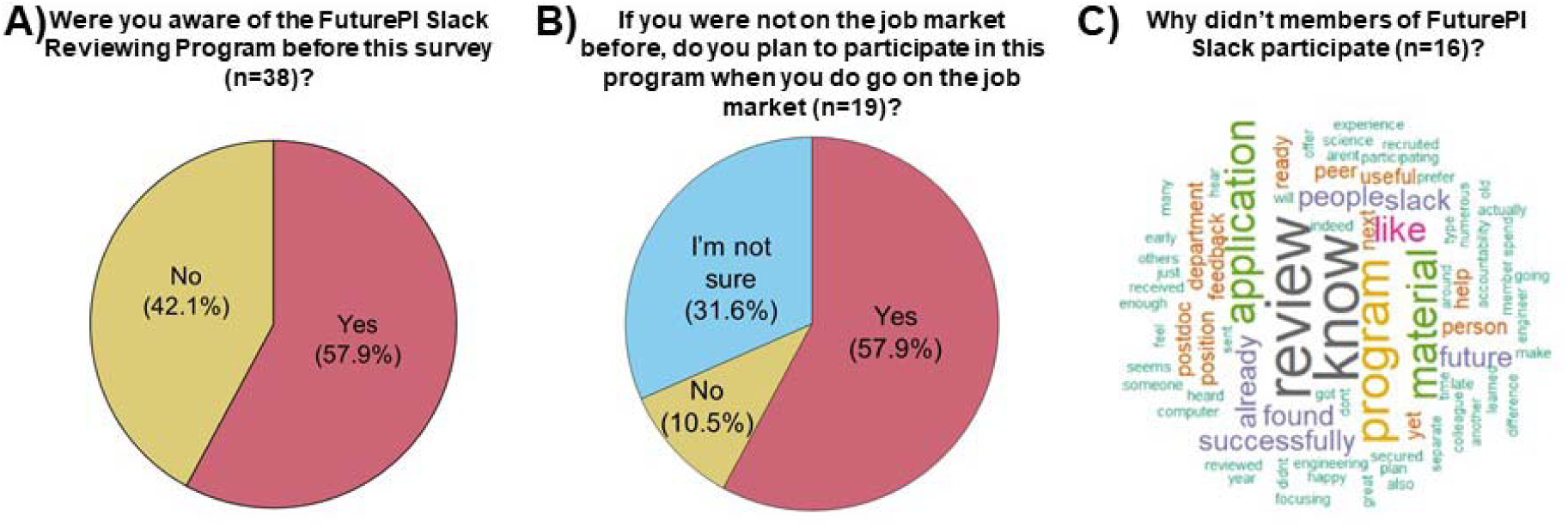
For non-participants, we asked if A) they were aware of the Program prior to the survey and, B) if they were not on the job market in the previous year, did they plan to participate in the program the following year. C) Word clouds highlighting the most common responses to the open-ended question of why non-participants did not participate in the Program.

> *“I would prefer feedback from people who have already successfully secured the type of position I would like*.*”*
>
> *“The main reason I did not think feedback from the peer reviews would be useful was that I assumed the majority of reviewers were peers who had yet to successfully get a job, and therefore their comments would be less useful than those of people who had successfully applied for and gotten an academic position, or people who had served on search committees. I do not know how this issue could be addressed within the Future PI Slack community, except to ask those who have since moved on to the next career stage to contribute back*.*”*
>
> *“I’d like to participate in this program if I know this review program before my applications*.*”*

## Discussion

For the past three years, we have been organizing a peer-reviewing program within FuturePI Slack for faculty job seekers. Our participants are diverse in their identities, geographic range, fields, current/training institutional type, and dream job type, a likely benefit of online organizing which does not limit us to recruiting participants from a specific location or institution. The results of the survey suggest that the Program benefits from the participants’ shared and unique characteristics and experiences both as a method for improving job application materials and as a significant mechanism of peer-support and mentorship. The survey data and personal communication with the organizers suggest that the participants found the Program to be overwhelmingly positive and would highly recommend the Program to others during their job search. This support is especially critical given the extremely stressful and frustrating nature of searching for a faculty job (Fernandes et al. 2020). Unfortunately, we are unable to comment on the ability of the Program to improve the likelihood of successfully obtaining a faculty position given the timing of our survey and the difficulty in tracking previous participants. Thus, we have no way of testing if participants are more or less likely to achieve a faculty position than non-participants. Nevertheless, the survey data provide valuable information for improving and expanding the Program and promoting its example to other institutions and organizations.

Previous studies have shown that peer review as a form of mentoring and support may be a critical mechanism for increasing the retention and success of early-career academics. Peer review offers not only mentoring and professional development but peer support in general also offers unique psycho-social benefits of emotional support from colleagues with shared experience and/or career stage (Cree-Green et al. 2020, Dickson et al. 2021). This unique benefit is particularly important for individuals from marginalized or under-represented groups in academia (Blackford 2018, Brommer & Eisen 2007, Eisen & Eaton 2017, Yadav & Seals 2019). Being a postdoc can be much more isolating compared to being a PhD student (Bruckmann & Sebestyen 2017) due to lack of cohesive cohort and fewer individuals at the same career level per lab. In fact, postdocs are the least likely to feel a sense of belonging across all stages of the scientific journey, from early graduate student to full professors, a feeling that is exacerbated for individuals from underrepresented or marginalized populations (Stachl & Beranger 2020). Data suggest, however, that having a scientific and social community play a significant role in individual success (Brommer & Eisen 2007, Yssldyk et al. 2019). Further, socio-emotional support and encouragement promote persistence in research careers for early career researchers while lack of these types of support is associated with disengagement from research and possibly attrition (Lambert et al. 2020, Pyhalto et al. 2017, Vekkaila et al. 2016). Social support can even combat, to a small degree, the negative psychological impact of sexism and racism within academia (Rodrigues et al. 2021).

Our survey data suggest that at least the Program, and likely FuturePI Slack more broadly, provides vital support to individuals facing greater difficulty in finding mentors for career support, either because of their career trajectory or identity. First, the respondents of this study are more likely to identify as members of historically underrepresented or marginalized groups, e.g., as female (although survey respondents overall are more likely to identify as female: Cull et al. 2005, Sax et al. 2003) or as non-white men, than the pool of postdocs in the USA, where the majority of the respondents are employed. This result is particularly notable given that, in biomedical sciences at least, men and women are equally represented in early career stages of PhDs and postdocs (NSF 2017), but women are significantly less likely than men to transition to an independent PI position (Lerchenmueller & Sorenson 2018), representing only 40% of assistant professors and 30% of associate professors (Jena et al. 2015). The transition from postdoc to independent PI is a major barrier for individuals from underrepresented minority populations (Bhalla 2019, Meyers et al. 2018). However, targeted interventions (e.g. peer review programs or workshops) can significantly increase postdoc confidence in their ability to apply to faculty jobs, which is predicted to increase persistence (Yadav & Seals 2019) at the career stage when these postdocs are most likely to ‘leak’ out of the pipeline. The demographics of our survey respondents likely also influenced the broad range of jobs our participants apply to: female and underrepresented minority postdocs are more likely to be interested in pursuing both research-intensive and teaching-intensive jobs, rather than only research-intensive jobs, and postdocs who are three or more years into their position are less committed to remaining in academia than postdocs in their first or second year (Lambert et al. 2020). Alternatively, postdocs who receive less mentoring from their primary supervisor are less likely to pursue an academic research career (Scaffidi & Berman 2012).

Further, participants were less likely to ask their PI or other postdocs to review their job application materials than the non-participants. This difference may be due to lack of support more broadly and/or because nearly 40% of our respondents switched fields between their PhD and their current position, a trend that is becoming increasingly common (Zeng et al. 2019), and thus may lack a broad network within their current field. Compared to non-participant respondents, the participants have spent more years on the job market, applied to more jobs, and a wider range of jobs, possibly reflecting a wider net cast by individuals searching for a position for longer. Lastly, the responses to the open-ended survey questions on the benefits of participating in the program confirmed that participants frequently received positive affirmation on the quality of their materials, which helped them to gain confidence and combat imposter syndrome while in the midst of the grueling search for a faculty job. Multiple respondents, in commenting on the difficulty and stress of searching for a faculty position, specifically mentioned that participating in the Program helped them to feel less alone (e.g., Cree-Green et al. 2020, Jaremka et al. 2020) and previous surveys have reported that postdocs found the process of applying for jobs to be easier when they had a strong network of support (Zimmerman 2018). The participants were further emotionally bolstered by the act of helping others to improve their materials and the idea that as a community, early-career academics all rise by helping each other. Community building as a career development strategy provides opportunities (Blackford 2018), especially for historically excluded groups, to build social capital and networks that may enhance their career opportunities (Alfred et al. 2019). Similar peer review programs could fulfill the same function by providing peer support and mentorship for early-career academics at their institutions, particularly for those who may lack that support elsewhere.

In addition to quantifying the utility of the Program, another goal of this study was, by combining both the survey and additional participant feedback sent directly to the organizers, to identify areas of potential improvement and best practices for participating. For example, this year we began recruiting participants for the program in June 2021 and organized the first week of peer-reviewing in July instead of waiting until August. Because the COVID-19 pandemic has highlighted the need for flexibility, we also transitioned from sending out materials on Monday and asking for feedback by Friday to asking that materials be sent out on Thursday with feedback due back on Monday. One challenge we have consistently encountered is ensuring full participation for and from everyone who signs up. Often, potential participants sign up immediately after the program announcement on FuturePI Slack, but weeks later, these individuals may find themselves with too much other work to participate. We now send an email the day before to everyone who has signed up requesting confirmation that they are still willing and able to participate before assigning groups. Even with this precaution, however, a few times a year even confirmed participants discover they are unable to participate. In these instances, we try to maintain flexibility for those participants and ensure the full benefits of participation for the remaining group members. We now ask participants to notify us if a group member failed to send out their materials or failed to send back feedback on the documents of their group members. If a participant fails to do either of these actions more than twice after confirming their desire to participate, they are removed from participation for the rest of the year in order to ensure other participants do not miss out on feedback from group members too often. So far, we have not had to remove a participant. We also ask to be notified of any unprofessional review comments and immediately bar individuals who provide such comments from participating in any future rounds of peer review (Silbiger & Stubler 2019).

Multiple participants suggested the following best practices guidelines, which we plan to include in future Program materials: 1) Participate early when documents are still quite rough in order to get ‘big picture’ or overall feedback on the ideas and organization; 2) Participate more than once to get a breadth of feedback from at least 4-6 peer reviewers, but on a timeline of every other week to allow for significant revision of early drafts; 3) Participate again ∼2 weeks before an important or specific deadline to get specialized feedback on nearly polished documents for a particular application.

Finally, the last goal of this study was to provide a model for other organizations wishing to develop similar peer-reviewing programs, in part because trainees’ perceived institutional support drives career search efficacy for postdocs (St. Clair et al. 2017). Already, the authors have received anecdotal reports from former participants organically replicating the goals and structure of this program in other organizations (e.g., Plant Postdoc Slack, Victoria University of Wellington Postdoctoral Society, and the NIH Office of Intramural Training and Education) for graduate students, postdocs, and other early-career academics. As indicated in our survey, although 70-80% of the respondents are employed at institutions having offices for professional development whose services might be of use to them, the majority of respondents were either unaware of any offering of editing services or sure that the offices did not offer such services. About 30-35% of respondents reported that these offices did offer such editing services, but only 10-15% of all respondents had taken advantage of this service. Moreover, the organization of similar peer-reviewing programs is not limited to universities and colleges; discipline-specific professional societies and organizations or broader national organizations like the National Postdoctoral Association could organize similar programs, either as a workshop at annual conferences or a multi-week program over the longer term. In fact, the sponsoring of such groups by professional scientific societies has been spontaneously suggested by postdocs in surveys as a way to improve their support of postdocs (Shaw et al. 2015). To lower the effort required to start a similar program, we include a ‘starter’ package of materials in the supplemental materials we routinely use. The focus can also be expanded to offer peer-review for the other aspects of the faculty application journey such as job talks or chalk talks (e.g., Henderson et al. 2016) or shifted to other types of academic documents like grant applications, course syllabi, or reappointment/tenure dossiers.

Ultimately, programs like the FuturePI Reviewing Groups Program provide an opportunity to improve the quality of one’s job application materials, which may, in turn, improve one’s odds of success in attaining an independent faculty position. However, programs that build peer-support and mentorship networks for early-career academics may also play a role in retaining and strengthening a diverse academic workforce despite the structural leaks in the pipeline. Our hope is that this Program description inspires other organizations to create similar programs to support vulnerable early-career academics in their search for independence.

## Supporting information

Supplemental Materials - Program Starter Pack

Supplemental Table 1 - Survey Questions

## Supplemental Material

Supplemental Material - ‘Starter Pack’ of example sign-up sheet, text describing the program, Confirmation email and assignment email.

Supplemental Table 1 - Survey Questions

## Author Contributions

KEG conceived of and designed the study, ET and AGT designed the survey questions with assistance from CG, AS, DK, EK, and KEG. CG, EK, AGT, and KEG analyzed the data. KEG drafted the manuscript with assistance from AS, AGT, and ET. All authors provided edits and approved the final manuscript.

## Acknowledgments

The authors would particularly like to thank FuturePI Slack for their participation and peer support, without which this program would not continue. Dr. Grogan would like to thank Dr. Doug James, who first inspired this program with his “Teaching Triangles” exercise when I was a PhD student in his Certificate in College Teaching program at Duke University.

