## Supplemental Materials - Program Starter Pack for "The power of peer networking for improving STEM faculty job applications: a successful pilot program"

**Organization of the FuturePI Reviewing Groups**

Here the authors provide a general outline of our FuturePI Reviewing Groups Program and tips for organizing a similar platform at your institution. The reader will find dates that worked for our Program, based on the timeline that most Universities and Research Institutions in the USA announce job opportunities, their deadlines, and the estimated time, in our own experience, a candidate needs to have a complete job application. Accordingly, this model may not be equally reproducible in other institutions around the world or for other purposes such as peer review of grant applications or mock interviews. However, the authors hope that this material might serve as a starting point for designing other Programs of peer-review of academic materials.

**Program Tasks & Personnel**

The tasks are simple: the program manager(s) should email each participant to confirm participation a given week, put together the groups (see below section *“Organizational Tips”*), and email assigned groups of people with instructions on how to share their material, what to do, and when to return the reviewed documents. The number of people required to coordinate these activities will depend on the frequency of review and the number of participants, but in general 1-2 people should be sufficient. We estimate that per week, it takes the program manager(s) 1 h of their time to confirm weekly participation and assign review groups. If programmatic changes are made or improvement surveys conducted after the annual program has ended, some additional time may be needed to make the changes or create and analyze the surveys.

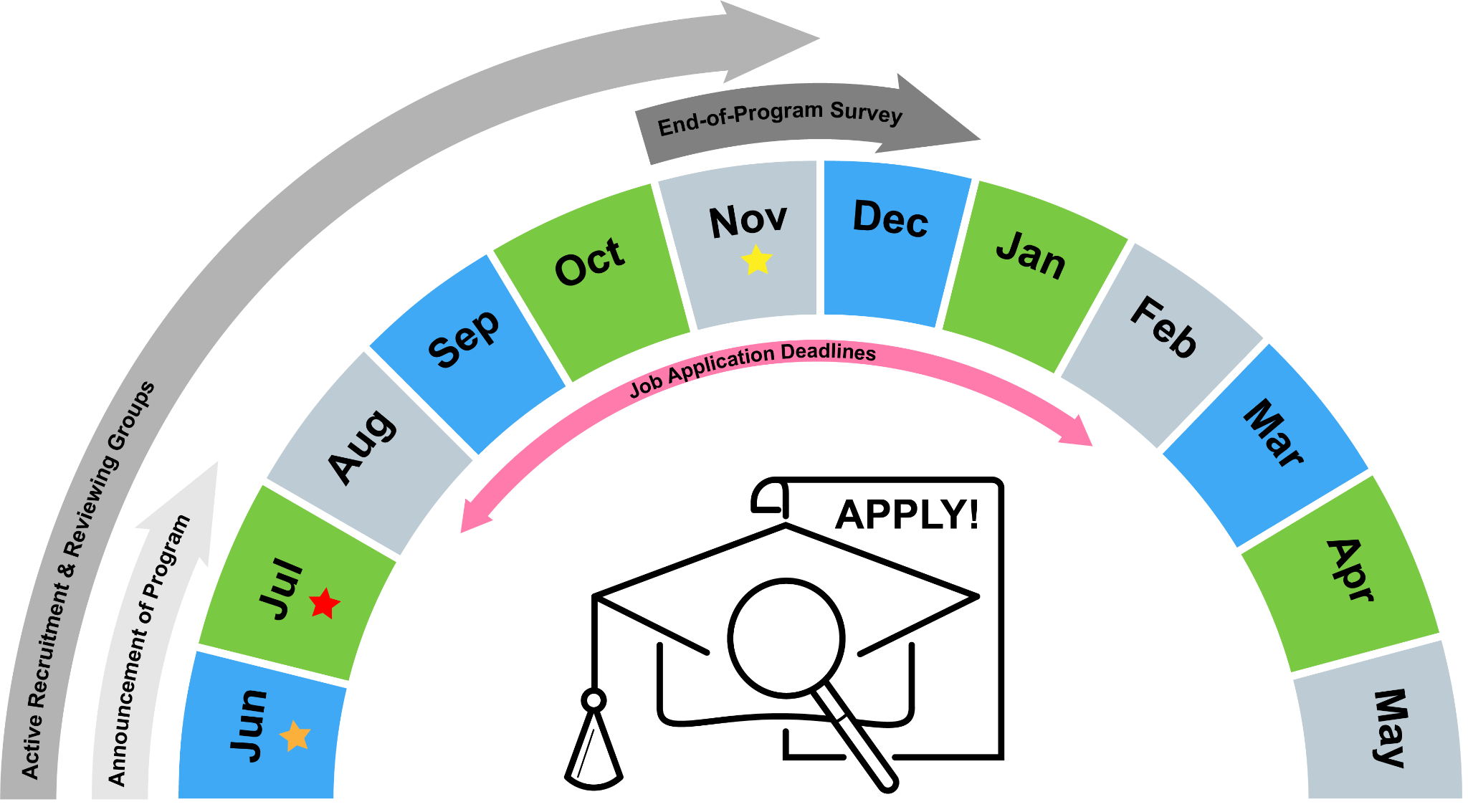

**Figure S1. Example Timeline used by the FuturePI Reviewing Groups Program.** *The academic job application season generally begins in August and continues through April of the following year, with the majority of job application deadlines occurring between September and January, although it is very common to find job posts with open deadlines and even late job advertisements at the beginning of the following year. To complement this timeline, the FuturePI Reviewing Groups Program starts announcements in June and continues during July to maximize its reach (orange star). During the same period, the sign-up goes live and the first reviewing groups are assigned in the middle of July (red star). Individuals can sign-up to participate in the Program at any time between June and November (yellow star), when the last groups are assigned.*

**Organizational Tips**

1. Send announcements explaining the goal and timeline of the program a month before the reviewing cycle starts (Fig. S1). For the FuturePI Reviewing Groups Program, we generally start announcing the program in June and the first reviewing groups are assigned in mid-July. Even if the applicant is not ready, knowing about the program could be a trigger to start preparing the application packages. An early announcement also gives a structure to those ready to apply and provides deadlines even before the job ads start appearing in the different platforms.
2. Clearly explain the goals of the program without the details and the logistics.
3. Use all the platforms available at your institution to advertise the program; we use Slack, but we widely announce the program through Twitter, Facebook, LinkedIn, email to colleagues and verbal communication.
4. 2 weeks before the starting date, send friendly reminders to boost the advertising of the opportunity.

***Example announcement email:***

*Good morning, FuturePI job seekers! The academic job season is about to start picking up and it's time to get your materials ready! We are kick-starting this year’s* ***Peer Reviewing Program*** *on the week of* ***July 12****. We use this channel to run a Peer Reviewing Program every Aug-Nov for job application materials. The purpose of this program is to provide both peer review support for those who may not have it and a broader audience to review your materials beyond the people in your lab or department; remember, search committees are often much broader in expertise than our own peer review networks.*

*Anyone who wishes to participate in this program can sign up here: [insert link of a sign-up sheet platform or request participants to send an email to the program manager]. You'll find the sign-up instructions at the top of the sheet. Provide your weekly availability, broad subject area, and your best email. Once we have this information, we will make groups of 3-4 members and email them with instructions to share their material and review it. The program works from* ***Monday to Friday every week,*** *and you can sign up as many times as you want your materials reviewed and you are willing to check other people’s. Usually, people sign up for 2-4 rounds of review, every two-three weeks, giving them multiple different reviewer audiences and in order to have time to incorporate all the changes between rounds.*

*Let’s get started! Looking forward to providing all the support to each of you job seekers out there. Good luck!*

5. For participant sign-up, we use a Google Sheet (Fig. S2) to collect all relevant information for contacting participants (name and best email address), as well as their field of study and the types of job they are applying to in order to best match them with peer reviewers. Additionally, the participants must indicate the week(s) they are willing to be part of the reviewing groups. That way, the program managers also can monitor the times each participant actively joined a group and avoid putting the same people in a group if they already were matched together in a previous week.
6. .

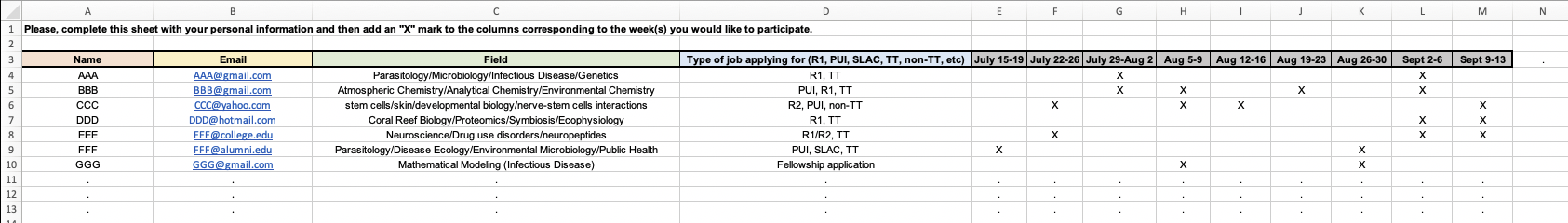

**Figure S2. Example of sign-up platform.** *Including information collected during sign-up, including name, email, field of study, target job type, and weeks of desired participation.*

7. Organizers can use a second Google Sheet to track how the groups have been made every week, the frequency of people participating (or not), any problems (and their solutions), templates for emails, etc.
8. The day before the Reviewing ‘week’ starts (for our scheme, the week starts on Monday), the program manager(s) should send an email to confirm the availability and willingness to participate of anyone who signed up for that week. This confirmation is to avoid including individuals who signed up weeks earlier and discovered that they cannot participate for that week, either because they do not have time, their materials are not ready, or any other reason.

***Example reminder email:***

*Hi all,*

*You've signed up to participate this week in the FuturePI Reviewing Groups and I need each of you to confirm to me by 9 am EST on Monday if you want to participate. Then, I'll send out the assignments. If you don't confirm by 9 am EST on Monday, I'll assume you're not interested this week, which is totally fine! Thanks & enjoy the rest of the week!*

9. Once the desire to participate in a particular week is confirmed, the program manager assigns each person to a group.
10. When organizing groups, the manager(s) use a few different criteria, such as field of study, type of job the participants plan to apply to, and previous composition of groups so as to avoid grouping the same participants repeatedly. The exact way these criteria are prioritized depends on the particular composition of the week’s participants.
11. After that, the program manager emails participants the group assignments, either via one email per group or a single email to all participants.

***Example assignment email:***

*Hi all,*

*Thanks for confirming your participation in the FuturePI Reviewing Groups this week! Below is the information for your Group, including their emails. Please send your materials (e.g., including but not limited to CV, Cover Letter, Research Statement, Teaching Statement, Diversity Statement) to each other today and return your feedback by Friday. FYI, the point of sending the materials back BY FRIDAY is so that authors can revise the materials in time for the next reviewing groups if they want to participate again soon. Feel free to send all the documents or just one, whatever you want help with! Remember, however, that your reviewers have busy lives too, so please don't send an overwhelming amount of material. It would also be helpful in your email to remind your reviewers of the types of institutions you want a job at and point them towards any sections of your job documents you'd like particular feedback. Thanks for participating and we hope it's informative/instructive/helpful. Also, if a member of the Group does not send their materials or comments in a timely manner, please let me know!*

| XXX YYY | |
| --- | --- |
| XXX | |
| AAA | |

12. Given that participants are exchanging personal documents with potential competitors, it is recommended to include a *Disclaimer* in order to clarify that program organizers are facilitating the exchange, but are not responsible for any issues that arise from the sharing of materials for peer review. Also, your institution may require special disclaimers to be included.

***Example disclaimer:***

*By participating in the FuturePI Reviewing Groups (the “Groups”), you acknowledge that the information and materials that you receive are the intellectual private property of others. The concepts, ideas, structure, figures, experimental research proposals, etc. are free to use and discuss only in the context of the Groups and for no other purpose. You shall not directly or indirectly disseminate or otherwise disclose, deliver, or make available to others any of the materials or concepts provided in the Groups. The appropriation of ideas or concepts discussed in the Groups for personal research use or grant applications by you is expressly prohibited; except in cases where the ideas or concepts are (a) demonstrably in the public domain, or (b) can be verified as having been independently developed by you prior to participation in the Groups. We, the organizers, take no responsibility if others do not adhere to these requirements.*

13. If organizers are interested in participant feedback or program improvement, you may wish to administer an end-of-program survey (see Supplemental Table 1 for example survey questions). The survey is best administered 1-3 weeks after the end of the program.
